# Body shape as a visual feature: evidence from spatially-global attentional modulation in human visual cortex

**DOI:** 10.1101/2021.11.17.469004

**Authors:** Sushrut Thorat, Marius V. Peelen

## Abstract

Feature-based attention modulates visual processing beyond the focus of spatial attention. Previous work has reported such spatially-global effects for low-level features such as color and orientation, as well as for faces. Here, using fMRI, we provide evidence for spatially-global attentional modulation for human bodies. Participants were cued to search for one of six object categories in two vertically-aligned images. Two additional, horizontally-aligned, images were simultaneously presented but were never task-relevant across three experimental sessions. Analyses time-locked to the objects presented in these task-irrelevant images revealed that responses evoked by body silhouettes were modulated by the participants’ top-down attentional set, becoming more body-selective when participants searched for bodies in the task-relevant images. These effects were observed both in univariate analyses of the body-selective cortex and in multivariate analyses of the object-selective visual cortex. Additional analyses showed that this modulation reflected response gain rather than a bias induced by the cues, and that it reflected enhancement of body responses rather than suppression of non-body responses. These findings provide evidence for a spatially-global attention mechanism for body shapes, supporting the rapid and parallel detection of conspecifics in our environment.

## Introduction

The capacity limits of the human visual system require selecting visual input for further processing and conscious access (Carrasco, 2011; Chun et al., 2011). One way to do this is to select specific locations of the visual field through spatial attention and eye movements. However, when searching for task-relevant objects in our environment, the location of these objects is typically not yet known. In this case, selection may operate at the level of visual features, using a selection mechanism termed feature-based attention (Maunsell and Treue, 2006). To be an effective selection mechanism, feature-based attention would need to operate in parallel across the whole or part of the visual field, in order to then guide spatial attention to the location of the target object (Wolfe, 1994). While this could be a plausible mechanism of attentional selection, it raises a core question: what are the features of feature-based attention?

At a neural level, it has been proposed that feature-based attention may be restricted to features to which sensory neurons are systematically tuned (Maunsell and Treue, 2006). Accordingly, the neural mechanisms of feature-based attention have been studied extensively with experiments involving low-level features for which such tuning has been established, such as the orientations of Gabor patches (Kamitani and Tong, 2005; Liu et al., 2007; Jehee et al., 2011) and the movement direction of random dot patterns (Treue and Trujillo, 1999; Saenz et al., 2002; Serences and Boynton, 2007). These experiments assessed how making one feature task-relevant influenced the responses of neurons that were selective or non-selective to that feature. A common finding was that attending to a low-level feature increased the responses of neurons selective to that feature and decreased the responses of neurons non-selective to that feature (Maunsell and Treue, 2006). Crucially, such modulations were shown to occur for stimuli presented in spatially-unattended and task-irrelevant locations (Treue and Trujillo, 1999; Saenz et al., 2002; Serences and Boynton, 2007; Zhang and Luck, 2009), providing evidence for a spatially-global mechanism of feature-based attention that can be distinguished from the effects of spatial attention.

In the present study, we tested whether global attentional modulation can similarly be observed for the shape of the human body, a category of high social and biological significance that is selectively represented in high-level visual cortex (Downing et al., 2001; Peelen & Downing, 2005). Behavioral studies have shown that bodies, like faces, gain preferential access to awareness (Stein et al., 2012) and automatically attract attention (Downing et al., 2004; Ro et al., 2007). There is also behavioral evidence for spatially-global attention effects for bodies: in a series of studies, spatial attention was captured by body silhouettes when participants searched for people in scenes presented in different parts of the visual field (Reeder and Peelen, 2013; Reeder et al., 2015). Finally, an fMRI study reported spatially-global modulation of multivoxel activity patterns distinguishing natural scenes with people from natural scenes with cars (Peelen et al., 2009). However, in that study, the relative contributions of scene context and of body, face, and car features could not be distinguished, such that it remains unknown whether feature-based attention effects exist for human bodies.

Here, we used fMRI to test for spatially-global attentional modulation of body processing in visual cortex. Participants detected the presence of bodies or one of five other categories (beds, bottles, cars, chairs, lamps) in task-relevant vertically-aligned images, thereby manipulating the top-down attentional set. To test for spatially-global attentional modulation, all analyses focused on responses evoked by objects that were concurrently presented at locations that were never relevant for the object detection task across three experimental sessions (Fig. 1A). The inclusion of five non-body categories reduced the possibility that participants could use a low-level feature to detect the presence of bodies, for example by looking for vertical (bodies) vs horizontal (e.g., cars) stimuli: lamps and bottles shared the vertical orientation with bodies (Fig 1D). To further reduce this possibility, each category was represented by a large and diverse set of exemplars cropped out of scene photographs. Finally, the use of silhouettes avoided possible low-level differences between categories in texture and color, and ensured that attention was guided by body shape rather than facial features (Störmer et al., 2019).

**Figure 1:**
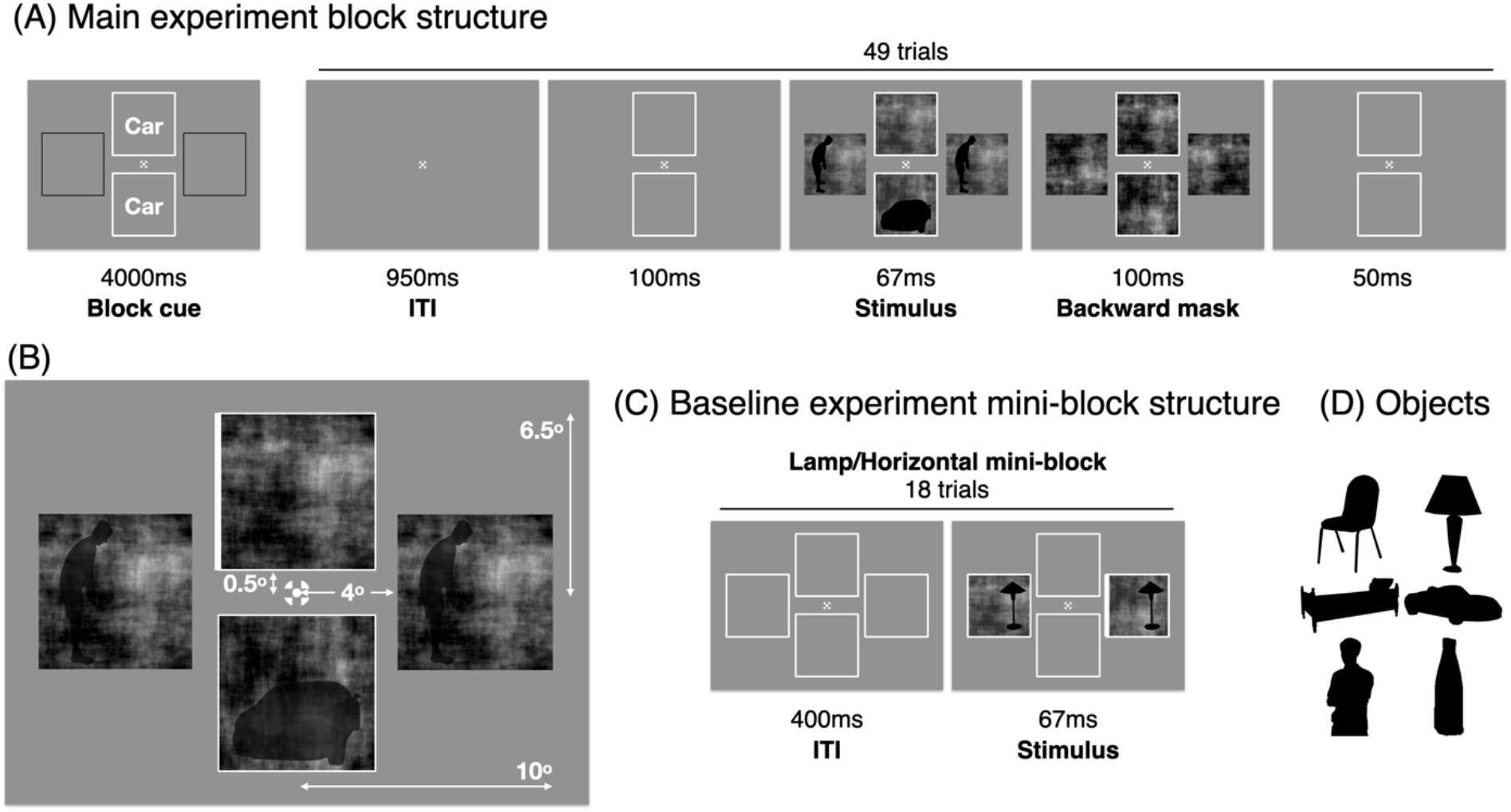
Experimental design. (A) The main experiment was designed to reveal the modulatory influence of feature-based attention on object responses evoked by stimuli presented at task-irrelevant locations (horizontal boxes). In each block (49 trials), participants had to search for the cued object category (e.g., car) in the vertical boxes, while objects were simultaneously presented in the horizontal boxes. (B) The spatial layout of the search display. (C) The baseline experiment was included to localize body-selective regions of interest (for the univariate analyses) and to obtain prototypical object category response patterns (for the multivariate analyses). Responses evoked by task-irrelevant objects in the main experiment were compared to these responses. Participants had to indicate if one of the edges of the two boxes thickened. The object category and location (horizontal or vertical boxes) varied across the mini-blocks. Unlike in the main experiment, the stimuli were not backward masked in order to increase visibility. (D) Exemplars of the six object categories: chairs, lamps, beds, cars, human bodies, bottles. Fifty exemplars were used for each category.

## Materials and Methods

### Participants

Twenty-three healthy adult volunteers with normal or corrected-to-normal vision gave written informed consent and participated in the experiment. All participants took part in three experimental sessions, on different days. One participant was excluded because of low performance on the visual search task (the difference between the proportion of false alarms and hits was lower than two standard deviations from the average difference). Twenty-two participants (mean age: 25.36 years; age range: 20-32 years; 11 female) were included in the reported analyses. The study was approved by the local ethics committee (CMO Arnhem-Nijmegen).

### Experimental Paradigm

In the main experiment, on each trial, the display contained two boxes in the horizontal and vertical locations (Fig. 1). The vertical boxes had a white bounding frame, signifying their relevance. Each of the four boxes contained a random image containing the average power spectrum of the objects from the six categories with random phases. Objects were mixed with these random images. On each trial, an exemplar from one of the six categories could be presented in one of the two vertical boxes (1/7 probability each) or no object would be presented (1/7 probability). Simultaneously, an exemplar from one of the six categories could be presented in both the horizontal boxes (1/7 probability each) or no object would be presented (1/7 probability). Each block consisted of 49 trials to fill the co-occurrence matrix of the horizontal and vertical object conditions, such that the conditions presented in the horizontal and vertical boxes were orthogonal to each other.

In each block of the main experiment, participants would either search for one of the six categories in the vertical boxes or would detect a thickening of the frames of the bounding boxes in the vertical location (for trial layout, see Fig. 1). Participants pressed the response button when the cued object category was shown in one of the vertical locations, which occurred on 7/49 trials. Participants had to respond within 1.2s. The brief presentation duration (67 ms) required participants to maintain fixation to be able to detect the target in one of the two vertical locations. Participants were instructed that they could ignore the objects presented at the horizontal locations. In the thickening condition, participants had to indicate, by pressing the response button, when one of the sides of the two bounding boxes became thicker than the others (thickening occurred on 7/49 trials in all blocks). Data from these thickening task blocks in the main experiment were not further analyzed because the within-block comparisons provided a more stringent test of our hypotheses, controlling for block-based effects (e.g., related to the processing of the category cue itself). The simultaneously presented objects in the horizontal boxes were always task-irrelevant. Each run contained four blocks, all containing a different search condition, such that across the seven search runs in each fMRI session each search block occurred four times. Feedback about search performance was provided at the end of each block.

In addition to the main experiment, participants completed a “baseline” experiment. This experiment was included to localize body-selective regions of interest (for the univariate analyses) and to obtain prototypical object category response patterns (for the multivariate analyses). In the baseline experiment, in different blocks, exemplars of one of the six categories or scrambled exemplars of one of the six categories were presented in both the boxes in either the horizontal or vertical locations (the other location left empty). These objects were mixed with a random image containing the average power spectrum of the objects from the six categories with random phases. The seven object conditions (six object categories and a scrambled objects condition containing a mix of scrambled objects from the six categories) and two presentation locations were blocked into mini-blocks containing 18 trials each. In each mini-block, participants had to search for thickening of the frames of the boxes where objects were being presented (1/7 probability of presence; each pair of thickening events had at least two non-thickening trials between them). Each block contained seven mini-blocks, with distinct object-location pairing, such that across the four blocks in each baseline experiment run, each type of block occurred twice. At the end of each block performance feedback was provided.

Each participant attended three experimental sessions. The first behavioral session required each participant to get exposed to the entire set of objects followed by the completion of one run of the baseline experiment and two runs of the main experiment. The second and the third sessions involved fMRI. In each of those sessions, the participant first browsed through the entire set of objects at their own pace, and then performed one run of the main experiment during the anatomical scan. This was followed by the functional recordings as the participants performed one run of the baseline experiment followed by four runs of the main experiment followed by one run of the baseline experiment followed by three runs of the main experiment.

### Stimuli

The stimulus presentation dimensions are shown in Fig. 1B. We acquired 50 exemplar silhouettes in real-world poses for each of the six categories of interest (beds, bottles, cars, chairs, lamps, and human bodies; shown in Fig. 1D). We obtained scenes containing the relevant objects from the SUN2012 database (Xiao et al., 2010) and Google images which were “Labelled for non-commercial reuse with modifications”, cropped out the objects, scaled them such than on one of the axes of the objects extended throughout the image, and converted them to silhouettes.

On each trial, the chosen exemplars were shown in the boxes, embedded in noise as mentioned above. The location of the objects within the boxes was jittered to increase variability. Objects that extended throughout the image horizontally were presented in one of three places within the box: touching the upper side, centered, or touching the lower side of the box. Similarly, objects that extended throughout the image vertically could be placed touching the left side, centered, or the right side of the box. The horizontally-placed boxes in the display contained the same stimulus (Fig. 1C).

### fMRI data acquisition and preprocessing

Functional (echo-planar imaging (EPI) sequence; 66 transverse slices per volume; resolution: 2×2×2mm; repetition time (TR): 1s; time to echo (TE): 35.2ms; flip angle: 60°; 6x multi-band acceleration factor) and anatomical (MPRAGE sequence; 192 sagittal slices; TR: 2.3s; TE: 3.03ms; flip angle: 8°; 1×1×1mm resolution; FOV: 256mm) images were acquired with a 3T MAGNETOM Skyra MR scanner (Siemens AG, Healthcare Sector, Erlangen, Germany) using a 32-channel head coil.

The functional data were analyzed using MATLAB (2017a) and SPM12. During preprocessing, within each session, the functional volumes were realigned, co-registered to the structural image, re-sampled to a 2×2×2mm grid, and spatially normalized to the Montreal Neurological Institute 305 template included in SPM12. Data were high-pass filtered with a cut-off of 128s. Temporal autocorrelations were accounted for using the AR(1) method in SPM. A gaussian filter (FWHM 3 mm) was applied to smooth the images.

### Statistical analysis

For each participant, general linear models (GLMs) were created to model the conditions in the experiment. In the main experiment, the GLM included regressors for the 49 conditions of interest: 7 attention blocks x 7 stimulus conditions presented in the task-irrelevant (horizontal) locations. As this was an event-related design, the onsets of the stimuli were modelled as impulse functions (delta functions) and the time series was convolved with the canonical HRF. In the baseline experiment, the GLM included regressors for the 14 conditions of interest: 7 stimulus conditions x 2 presentation locations. As this was a block-design, the mini-blocks corresponding to each stimulus condition were modelled as boxcars and the time series was convolved with the canonical HRF. Separate GLMs were executed for each run of the main and baseline experiments. The acquired regression weights were averaged across repetitions of the corresponding conditions across the runs. Regressors of no interest were also included to account for differences in the mean MR signal across scans and for head motion within scans.

In the univariate analysis, the regression weights (betas) from the GLM were compared between conditions after averaging across the voxels of a region of interest (ROI). In the multivariate analysis, the pattern of betas from the GLM across the voxels of an ROI were compared between conditions using Kendall’s tau correlation coefficient (τ) as a metric for similarity. Before comparing the betas between the main and baseline experiments, the data were mean-centered: the mean across all main experiment condition betas was subtracted from those condition betas (separately for each voxel), and the mean across all baseline experiment condition betas were subtracted from those condition betas.

### Regions of interest

All ROIs were defined across both hemispheres (except FBA, which was limited to the right hemisphere). In the multivariate analysis, we focused on two ROIs, the lateral-occipital cortex (LOC) and the early visual cortex (EVC). The LOC ROI was defined using a group-constrained subject-specific method (Fedorenko et al., 2010). The group-level ROI was defined by first contrasting the average response to the 6 object categories with the response to the scrambled objects in the baseline experiment. Threshold-free cluster enhancement (TFCE; Smith and Nichols, 2009) with a permutation test was used to correct for multiple comparisons (at p < 0.05) across the whole brain. The resulting voxels were intersected with the lateral occipital cortex ROI from Julian et al. (Julian et al., 2012) to obtain the group-level LOC ROI. Then, for each participant, the 1000 most object-selective voxels (average object response - scrambled stimulus response, in the baseline experiment horizontal conditions) within the group-level LOC ROI were selected for further analysis. The EVC ROI was defined at the individual participant level as the 1000 most responsive voxels (average object response > 0, in the baseline experiment horizontal conditions) in Brodmann area 17 (corresponding to V1; Wohlschläger et al., 2005). Brodmann area 17 was taken from the Brodmann atlas available in SPM12.

In the univariate analysis we focused on two body-selective ROIs, the extrastriate body area (EBA; Downing et al., 2001) and the fusiform body area (FBA; Peelen and Downing, 2005). The ROIs were defined using the method described above for LOC. The group-level ROI was defined by first contrasting the response to bodies with the average response to the other 5 categories in the baseline experiment. TFCE was used to correct for multiple comparisons (at p < 0.05) across the whole brain. The resulting voxels were intersected with ROIs from Julian et al. (2012): the extrastriate body area ROI to obtain the group-level EBA ROI and the fusiform face area (FFA) ROI to obtain the group-level FBA ROI (FBA is not provided, but the FFA and FBA closely overlap at the group-level; Peelen and Downing, 2005). Then, for each participant, the 20 most body-selective voxels (body response - average response to other objects, in the baseline experiment horizontal conditions) within the group-level ROIs were selected for further analysis.

### Multivariate analysis approach

In the multivariate analyses, we correlated multivoxel activity patterns evoked by the task-irrelevant objects in the main experiment with multivoxel activity patterns evoked by the clearly visible objects in the baseline experiment, using Kendall rank-ordered correlation; *τ*. We expect to find stronger correlations between corresponding object categories (e.g., between bodies in the main experiment and bodies in the baseline experiment), than between non-corresponding categories (e.g. between bodies in the main experiment and beds in the baseline experiment). As such, the difference between corresponding and non-corresponding category correlations is a measure of category processing (Peelen et al., 2009), analogous to decoding accuracy. Here, we computed *proximity* to the categories in the baseline experiment as the correlation with that category minus the correlation with the other categories in the baseline experiment. For example, for bodies, the proximity to bodies (in the baseline experiment) is the correlation between bodies in the main experiment and bodies in the baseline experiment minus the average correlation between bodies in the main experiment and the other five categories in the baseline experiment.

### Image-based discriminability approach

To rule out that bodies differed systematically from the other objects in terms of low-level features, we used representations of the exemplars in the layers of a convolutional neural network (trained for object recognition in natural images; CNN; AlexNet: Krizhevsky et al., 2012) to test for image-based categorizability differences across the categories. Output activations at each layer corresponding to 50 exemplars of each of the six categories, embedded in noise as in the fMRI experiment, in the three possible locations defined by the shapes (see the subsection on Stimuli), were extracted. Balanced linear support vector machines (SVM) were trained to classify between the images of one category (150 images each) as opposed to the other categories. 10-fold cross-validated classification accuracies were reported for each category for each layer of the CNN.

## Results

In the main experiment, participants detected the presence of object silhouettes belonging to one of six categories (Fig. 1D), in different blocks. Throughout the experiment, only the vertically-aligned locations were relevant for the detection task (Fig. 1A). Each block started with a category cue (e.g. “Car”) indicating the target category for that block (Fig. 1A), followed by 49 object detection trials. In 42 trials (6/7th), one of the two task-relevant locations contained a briefly-presented object (67 ms) within phase-scrambled noise (Fig. 1B), with each category presented equally often (7 trials each). In the remaining 7 trials (1/7th) no object was presented.

Crucially, in 6/7th of the trials, two objects were simultaneously presented in the horizontally-aligned locations (Fig. 1A). These objects were briefly presented (67 ms), embedded in noise, and backward masked. This was done to reduce the possibility of participants moving their eyes (and/or spatial attention) to the task-irrelevant objects. Objects at the horizontal locations were never relevant for the participants and participants were instructed that these could be completely ignored. The occurrence probabilities of the categories were the same as for the task-relevant locations. The 7 vertical and 7 horizontal conditions were fully crossed within each block, resulting in 49 trials, which were presented in random order. Trials were coded according to the categories presented in the horizontally aligned (task-irrelevant) locations, as these were the focus of our analyses.

### Task performance (task-relevant locations)

Averaged across the two fMRI sessions and across object search blocks, participants had a hit rate of 78.3% (proportion of the target-present trials where participants responded) and a false alarm rate of 5.6% (proportion of the target-absent trials where participants responded), resulting in an average d’ (zscore(hit rate) - zscore(false alarm rate)) of 2.7 (beds: 2.0; cars: 2.4; bottles: 2.6; bodies: 2.9; chairs: 2.9; lamps: 3.3). Note that this was the performance for the task-relevant stimuli presented at the vertical locations. All fMRI analyses focused on the objects presented at the task-irrelevant horizontal locations. For responses to objects at the task-irrelevant locations, we refer the reader to the section *The relationship between attentional modulation and behavioral responses*.

### Univariate results in EBA and FBA

Previous research has shown that bodies evoke a selective univariate response in two focal regions of high-level visual cortex: the extrastriate body area (EBA; Downing et al., 2001) and the fusiform body area (FBA; Peelen & Downing, 2005). Here, EBA and FBA were defined based on responses in the baseline experiment (see Material and Methods). We tested for spatially-global attention effects for bodies in these ROIs by comparing body-selective responses in EBA and FBA evoked by task-irrelevant bodies across target detection blocks in the main experiment. Betas were averaged across the voxels of each ROI to acquire one beta per condition for each ROI. For each category, the beta corresponding to within-block trials in which no objects were presented was subtracted to account for block effects. Responses to non-body objects and non-body detection blocks were averaged, such that we had 4 values for each ROI: body and non-body stimuli, presented in the body and non-body detection blocks. The difference between body and non-body stimuli within each block is a measure of body selectivity.

A 2 (ROI) x 2 (attention: body, other categories) repeated-measures ANOVA on body selectivity (response to bodies minus average response to other categories) revealed a main effect of attention (F_1,21_ = 7.2, p = 0.014, η^2^_p_ = 0.25), reflecting stronger body selectivity in body attention blocks than non-body attention blocks (Fig. 2A). This attention effect interacted with ROI (F_1,21_ = 4.6, p = 0.043, η^2^_p_ = 0.18), being stronger for EBA than FBA. When analyzed separately, both EBA and FBA showed an attention effect, such that body selectivity was higher in body detection blocks than in other category detection blocks (EBA: F_1,21_ = 7.4, p = 0.013, η^2^_p_ = 0.26; FBA: F_1,21_ = 4.4, p = 0.049, η^2^_p_ = 0.17; Fig. 2A). The attention effect was consistent across ROI sizes (Fig. 5). In EBA, bodies evoked a selective response in both the body detection blocks (t_21_ = 4.6, p < 0.001, d = 1.00) and the other category detection blocks (t_21_ = 4.4, p < 0.001, d = 0.96), while in FBA body selectivity was only positive in the body detection blocks (t_21_ = 2.5, p = 0.02, d = 0.55; other category detection blocks: t_21_ = 0.8, p = 0.42, d = 0.17).

**Figure 2:**
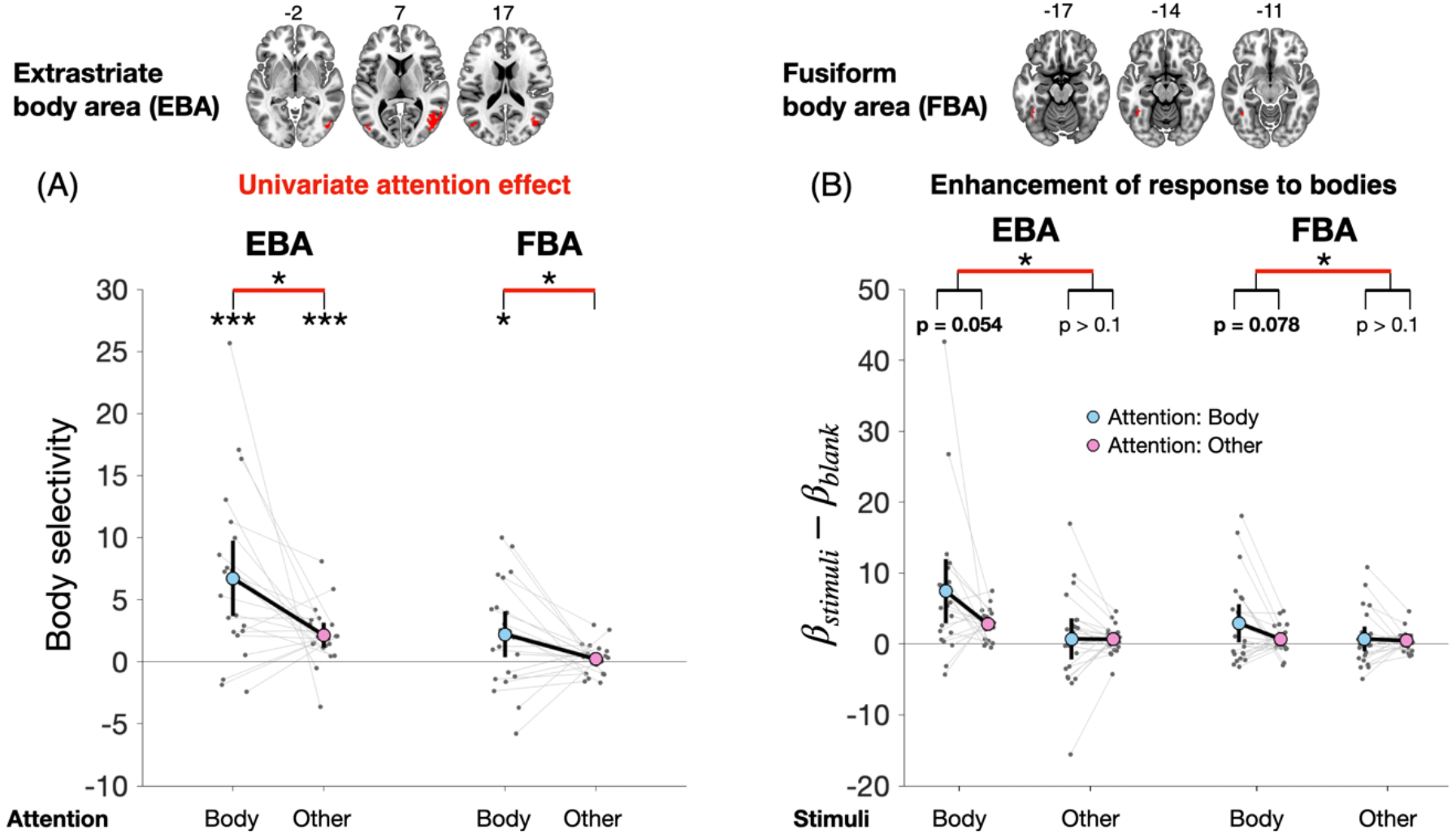
Univariate attention effect in body-selective ROIs. (A) Body selectivity (response to body - average response to other objects) was higher when bodies were attended, in both ROIs. This provides evidence for spatially-global attentional modulation for body silhouettes. (B) Across ROIs, the response to bodies (corrected for block-wise differences by subtracting the corresponding blank responses) was enhanced while the responses to other categories remained unchanged. Error bars indicate 95% confidence intervals for the measures indicated on the y-axes. The asterisks indicate p-values for the t-tests of the corresponding comparisons (*p< 0.05, **p<0.01, ***p< 0.001). EBA and FBA ROIs are displayed together with MNI z-coordinates.

The attention effect for bodies in EBA and FBA could reflect enhanced responses to bodies presented in body detection blocks, but may also (or additionally) reflect reduced responses (suppression) to the other categories presented in body detection blocks. To test these alternatives, we compared body and object-evoked responses across the body and object-detection blocks (after subtracting the response to blanks within each block). Averaged across ROIs, there was a higher response to bodies in body detection blocks than in other category detection blocks, which was marginally significant (paired t-test, t_21_ = 2.1, p = 0.05, d = 0.46; Fig. 2B). There was no evidence that the response to the other objects was suppressed, with equally strong responses in both blocks (paired t-test, t_21_ = 0.19, p = 0.85, d = 0.04). These effects were also observed, though weaker, in each ROI separately (statistics provided in Fig. 2B).

These results provide the first evidence for spatially-global attentional modulation for body silhouettes, show that these effects are strongest in EBA, and link these effects to enhancement of body responses rather than suppression of non-body responses.

### Multivariate results in LOC

Previous studies have shown that multivoxel activity patterns in object-selective cortex distinguish between object shapes (Haushofer et al., 2008; Op de Beeck et al., 2008; Eger et al., 2008). This gave us another opportunity to test for spatially-global effects of attention, including for non-body categories. Here, instead of body selectivity, we used proximity (Pr) as dependent measure. Proximity was based on correlations between response patterns in the main experiment and response patterns in the baseline experiment, following previous work (Peelen et al., 2009). Proximity reflects how similar a category’s response pattern in the main experiment is to a category’s response pattern in the baseline experiment, relative to the other categories in the baseline experiment (Materials and Methods). For example, for bodies, the proximity to bodies (in the baseline experiment) is the correlation between bodies in the main experiment and bodies in the baseline experiment minus the average correlation between bodies in the main experiment and the other five categories in the baseline experiment.

### Attentional modulation for bodies in LOC

The proximity to bodies is shown in Fig. 3A. A 2 (attention: body, other categories) x 2 (stimulus presented: body, other categories) repeated-measures ANOVA revealed a significant interaction (F_1,21_ = 30.4, p < 0.001, η^2^_p_ = 0.59), reflecting a stronger difference between the proximities for body and non-body categories when participants attended to bodies (t_21_ = 9.9, p < 0.001, d = 2.2) than when they attended to the other categories (t_21_ = 8.1, p < 0.001, d = 1.8). The multivariate attention effect in LOC was consistent across ROI sizes (Fig. 5). These results provide further evidence for spatially-global attentional modulation of body processing.

**Figure 3:**
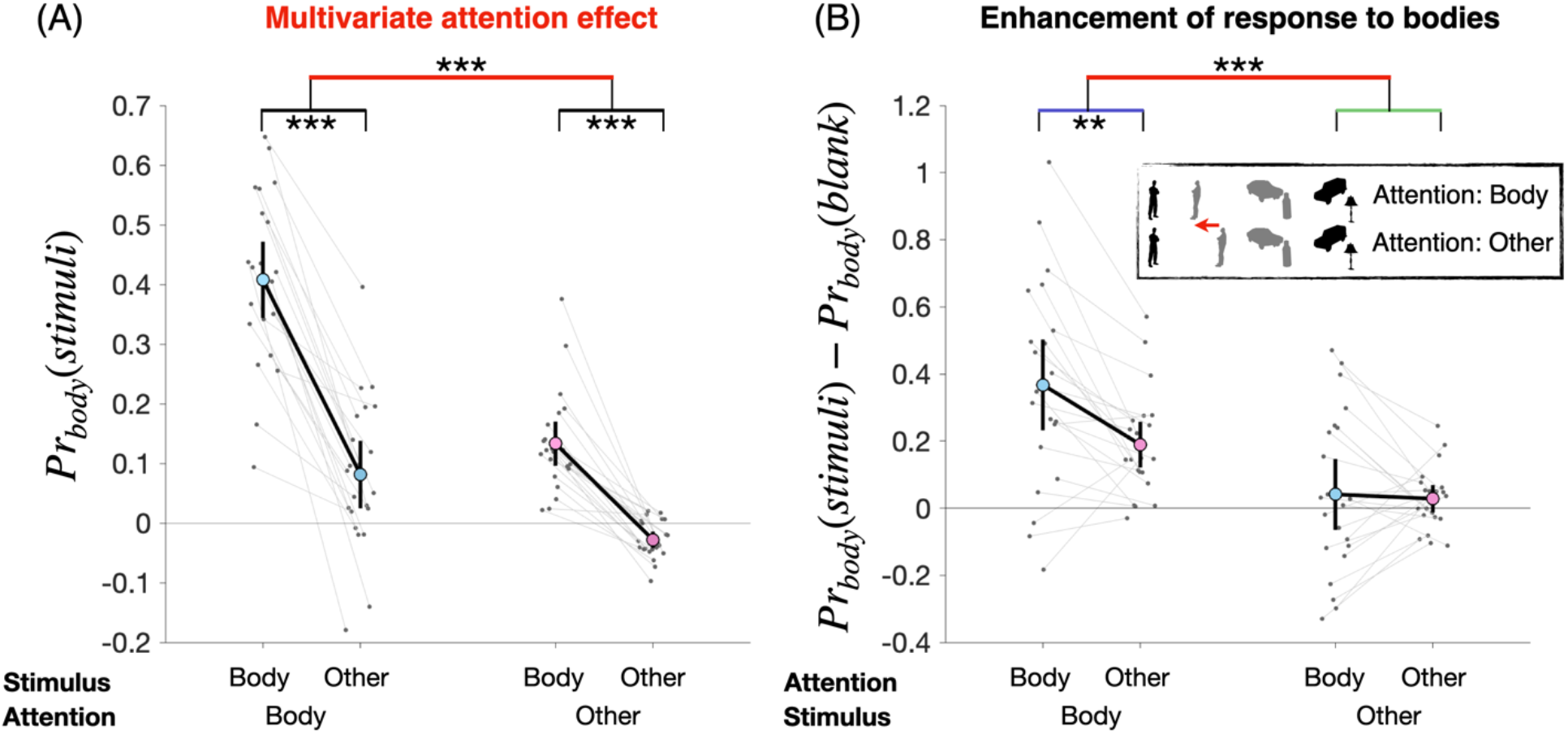
Probing the multivariate attention effect for bodies in LOC. (A) The selective proximity for bodies (proximity to bodies for Body vs Other) is higher when bodies are attended, which is evidence for a multivariate attention effect in LOC (comparison highlighted in red), reflecting response gain. (B) Proximity (to bodies) of bodies and other categories were compared between the body attention blocks and the other category attention blocks, corrected for block-wise differences by subtracting the proximity (to bodies) to blank responses within blocks. When bodies were attended, the proximity of bodies was enhanced, whereas the proximity of the other categories was not affected (inset: gray objects correspond to attention-dependent representations and black to benchmark representations). This indicated that the multivariate attention effect for bodies in LOC (the comparison corresponding to the red bar) was driven primarily by enhancement of body-selective response patterns when bodies were attended. 95% confidence intervals for the measures indicated on the y-axes are shown. The asterisks indicate the p-values for the t-tests of the corresponding comparisons (*p< 0.05, **p<0.01, ***p< 0.001). Blue: attentional modulation for bodies; green: attentional modulation for other categories.

The attention effect for bodies in LOC could reflect enhanced proximity to bodies for the bodies presented in body detection blocks, but may also (or additionally) reflect reduced proximity to bodies (suppression) for the other categories presented in body detection blocks. To test for body-selective enhancement, we compared the proximity (to bodies in the baseline experiment) for bodies in the body detection blocks with the corresponding proximity of other objects in the body detection blocks. To account for overall differences between blocks (e.g., related to the cue or to block-based attentional bias), we subtracted the proximity to bodies for the within-block trials in which no objects were presented. Results showed that proximity to bodies was significantly enhanced for bodies presented in the body detection blocks as compared with bodies presented in the other detection blocks (t_21_ = 3.5, p = 0.002, d = 0.76; blue comparison in Fig. 3B). There was no evidence for suppression: proximity to bodies was not different for objects presented in the body detection blocks as compared with objects presented in the other detection blocks (t_21_ = 0.3, p = 0.78, d = 0.06; green comparison in Fig. 3B). The difference between these effects (red comparison in Fig. 3B) corresponds to the same multivariate attention effect as shown in Fig. 3A. These results show that the multivariate attention effect was primarily driven by the enhancement of body-selective response patterns, in line with the univariate results (Fig. 2).

### The relationship between attentional modulation and univariate body selectivity of LOC voxels

Next, we tested whether the multivariate attention effect observed for bodies in LOC depended on the (univariate) body-selectivity of voxels included in LOC. To this end, we computed the multivariate attention effect for bodies in an ROI that consisted of LOC voxels that responded less strongly to bodies than to other categories in the baseline experiment (on average 330.8 out of the original 1000 voxels satisfied this criterion). Results were compared with a size-matched ROI consisting of randomly-sampled LOC voxels (size-matching done within each participant; sampled 100 times). Attentional modulation was computed in the same way as for the whole LOC in the original analysis (red comparison in Fig. 3). Attentional modulation was stronger for the size-matched ROI than the non-selective ROI (t_21_ = 3.1, p = 0.006, d = 0.68). However, attentional modulation was significant even in the non-selective ROI (t_21_ = 2.1, p = 0.047, d = 0.46). These results suggest that the attentional modulation in LOC was partly but not exclusively driven by body-selective voxels.

### Attentional modulation for non-body categories in LOC

Using the multivariate analysis framework outlined above for bodies, we can similarly test for spatially-global attentional modulation for the other categories. For each non-body category, we computed the multivariate attention effect as was done for bodies, now using the proximity to that category in the baseline experiment. To reduce the complexity of the ANOVA and the corresponding visualization of the data, we used *selective proximity* as the dependent measure. Selective proximity is the proximity difference between the corresponding and non-corresponding categories (e.g., the difference between the two left-most data points in Fig. 3A). As an intuition for what this new measure represents, note that in the case of bodies, selective proximity is analogous to the body selectivity measure in the univariate analysis.

In LOC, a 6 (category of interest) x 2 (category attended/unattended) repeated-measures ANOVA on these selective proximities revealed a significant interaction (F_5,105_ = 3.9, p = 0.003, η^2^_p_ = 0.16; Fig. 3A), indicating that attention differentially affected the selective proximity of the six categories. Six paired-sample t-tests showed that attentional modulation was significant for bodies (t_21_ = 5.5, p_bonf_ < 0.001, d = 1.2; red comparison in Fig. 4A), as already shown in the previous analyses (Fig. 3). No significant multivariate attention effect was observed for the other categories (t_21_ < 2.4, p_bonf_ > 0.1, d < 0.5; for all tests; Fig. 4A).

**Figure 4:**
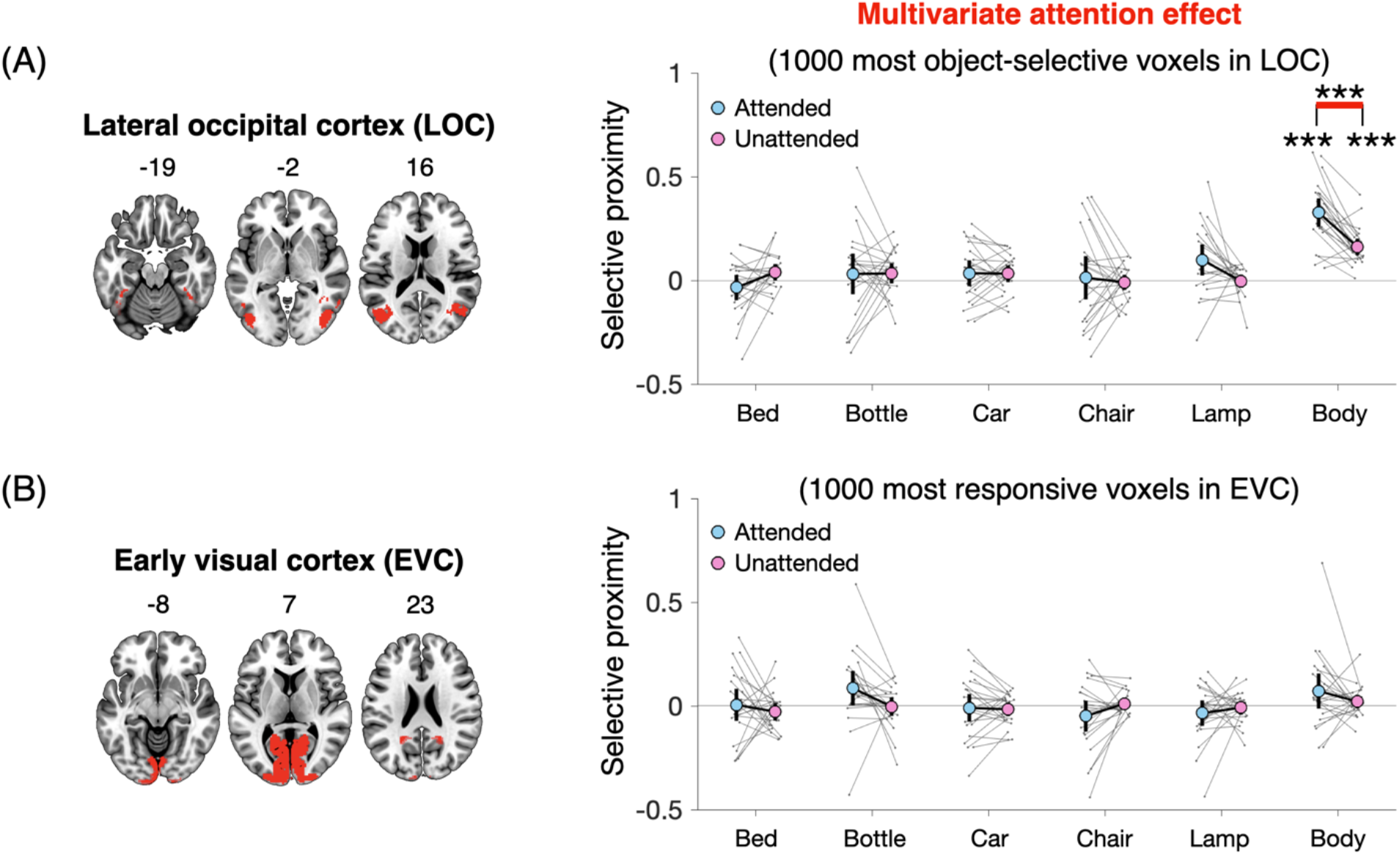
Multivariate attention effect. The selective proximities, for the attended and unattended conditions, are shown for the six categories in the two ROIs. The multivariate attention effect is the difference between attended and unattended selective proximity (comparison highlighted in red). A) In LOC, we find evidence for attentional modulation of the selective proximities of bodies. B) No attentional modulation was found in EVC. Error bars indicate 95% confidence intervals for the selective proximities. The asterisks denote Bonferroni corrected p-values for the t-tests of the twelve comparisons related to selective proximities, and Bonferroni corrected p-values for the t-tests of the six comparisons related to selective proximity modulations (*p< 0.05, **p<0.01, ***p< 0.001).

### Attentional modulation in EVC

The same analysis was conducted in early visual cortex (EVC; see Materials and Methods). A 6 (category of interest) x 2 (category attended/unattended) repeated-measures ANOVA on selective proximities revealed a marginally significant interaction (F_5,105_ = 2.2, p = 0.06, η^2^_p_ = 0.095; Fig. 4B), no significant main effect of attention (F_1,21_ = 0.6, p = 0.4, η^2^_p_ = 0.028), and a marginally significant main effect of category (F_5,105_ = 2.2, p = 0.06, η^2^_p_ = 0.096). Paired-sample t-tests showed no significant attentional modulation for any of the categories (|t_21_| < 2.2, pbonf > 0.1, d < 0.48; for all tests). Finally, attentional modulation for bodies was significantly stronger in LOC than in EVC (t_21_ = 2.9, p = 0.01, d = 0.63).

### The relationship between attentional modulation and behavioral responses

In both multivariate and univariate analyses, we found that the body-selective response elicited by body silhouettes in task-irrelevant locations was enhanced in body detection blocks compared with other category detection blocks. This raises the question of whether this attentional modulation affected behavior in the detection task. Particularly, did participants disproportionally false alarm to the bodies at task-irrelevant locations when detecting bodies at task-relevant locations? Because of the orthogonal design, each category (+blank stimulus) in the irrelevant location appeared equally often with each category (+blank stimulus) in the relevant location. Therefore, when the target category (e.g., bodies) appeared at the task-irrelevant location no target was presented at the task-relevant locations in most trials (6/7th), and participants had to withhold their response. For these trials, we tested whether responses (i.e., false alarms) depended on the combination of the category presented and the category that was the target in that block. To this end, for each category, we computed the difference between the false alarm rate (FA) to that category and the average FA to the other categories, separately for each block. We then compared this ΔFA for trials in which the object matched the target category (e.g., bodies presented in body blocks) and trials in which the object mismatched the target category (e.g., bodies presented in bed blocks).

A 2 (matching, non-matching) x 6 (target category) repeated-measures ANOVA on ΔFA revealed a significant interaction (F_5,105_ = 3.3, p = 0.008, η^2^_p_ = 0.14; Fig. 6). Six paired-sample t-tests showed that ΔFA was stronger when the object matched the target category for all categories (t_21_ > 2.9, p_bonf_ < 0.05, d > 0.63, for all non-body categories, biggest difference of 6.5% for cars; bodies: t_21_ = 2.79, p_bonf_ = 0.066, difference of 3.7%, d = 0.61). These results show that participants disproportionally false alarmed when the target category was shown at the task-irrelevant location. Contrary to the fMRI results, however, this effect was relatively weak for bodies.

**Figure 5:**
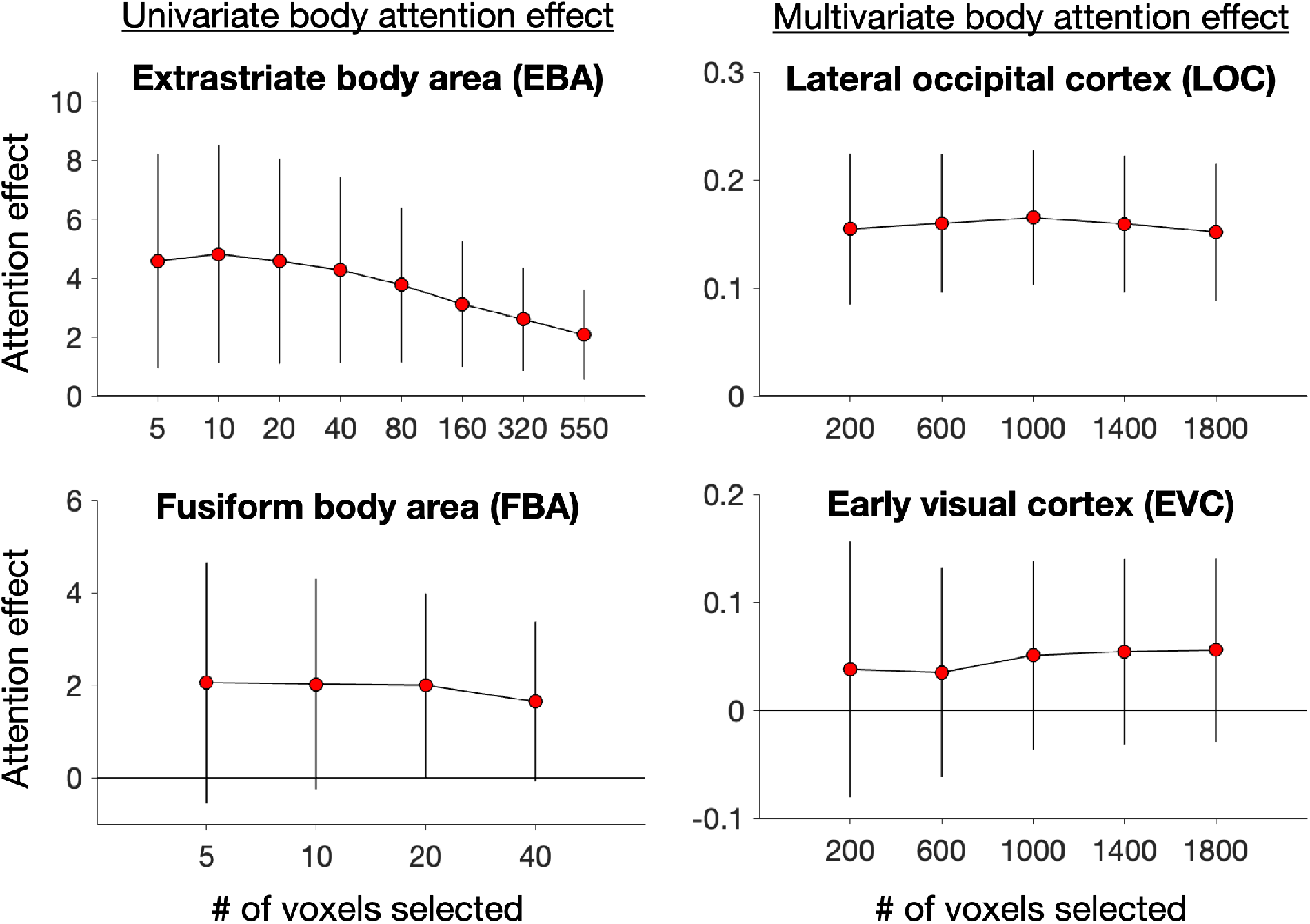
The influence of voxel selection on body attention effects. The univariate attention effects, for EBA and FBA, and the multivariate attention effects, for LOC and EVC, for bodies, are shown as a function of the number of voxels selected within each ROI. The attention effects for bodies observed in EBA, FBA, and LOC, are observed regardless of the number of voxels selected. Error bars indicate 95% confidence intervals for the attention effects.

**Figure 6:**
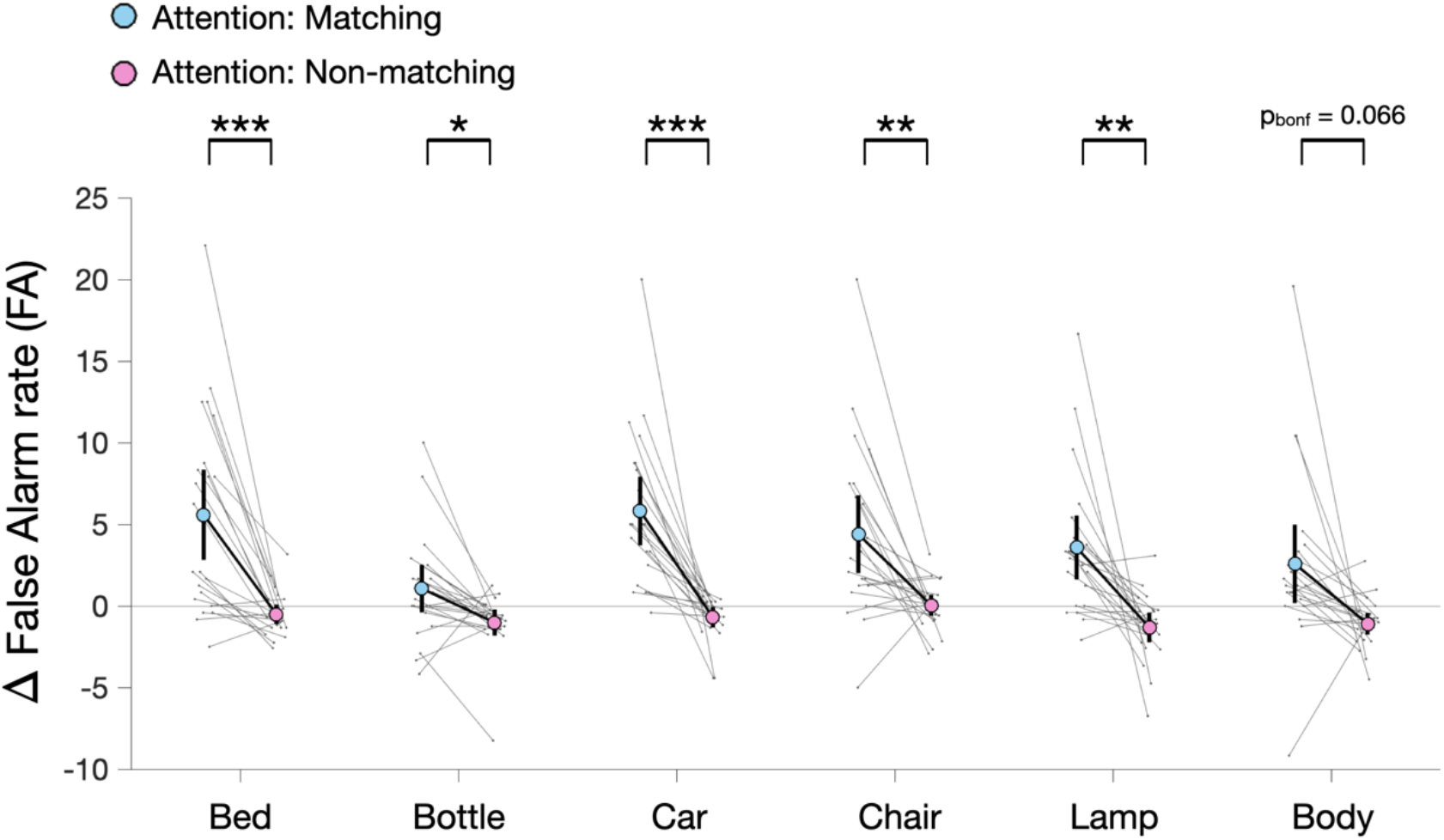
The relationship between attentional modulation and behavioral responses. Participants disproportionally false alarmed when the target category was shown at the task-irrelevant location (matching>non-matching) but this effect was relatively weak for bodies. Error bars indicate 95% confidence intervals for ΔFA. The asterisks denote instances where t-tests returned p_bonf_ < 0.05 for the corresponding comparisons (*p< 0.05, **p<0.01, ***p<0.001).

### Image-based discriminability

In all fMRI analyses, we found that bodies were more strongly represented and more strongly modulated by attention than the other categories. This could reflect an interesting property of bodies, for example, related to the life-time relevance of detecting conspecifics or to the increased familiarity with body shapes. However, it could potentially also reflect uncontrolled image-based differences: perhaps the body silhouettes included in the study stood out from the other objects in terms of low-level features. To exclude this possibility, we decoded object categories from the object exemplar representations in the layers of a convolutional neural network trained for object recognition (Materials and Methods). For each of the 6 categories, in each layer of the CNN, one-vs-all linear discriminant classifiers were trained to discriminate each category from the other categories using the 50 exemplars of each category presented in the fMRI experiment. 10-fold cross-validation accuracies were analyzed across the objects.

As shown in Fig. 7, bodies were less discriminable than most other categories in the early layers of the CNN. It is only in the mid to final layers - where overall classification is almost at ceiling - that the classification accuracy for bodies is similar to the average accuracies for the other categories. This result shows that the image-based discriminability was, if anything, lower for bodies than for the other objects.

**Figure 7:**
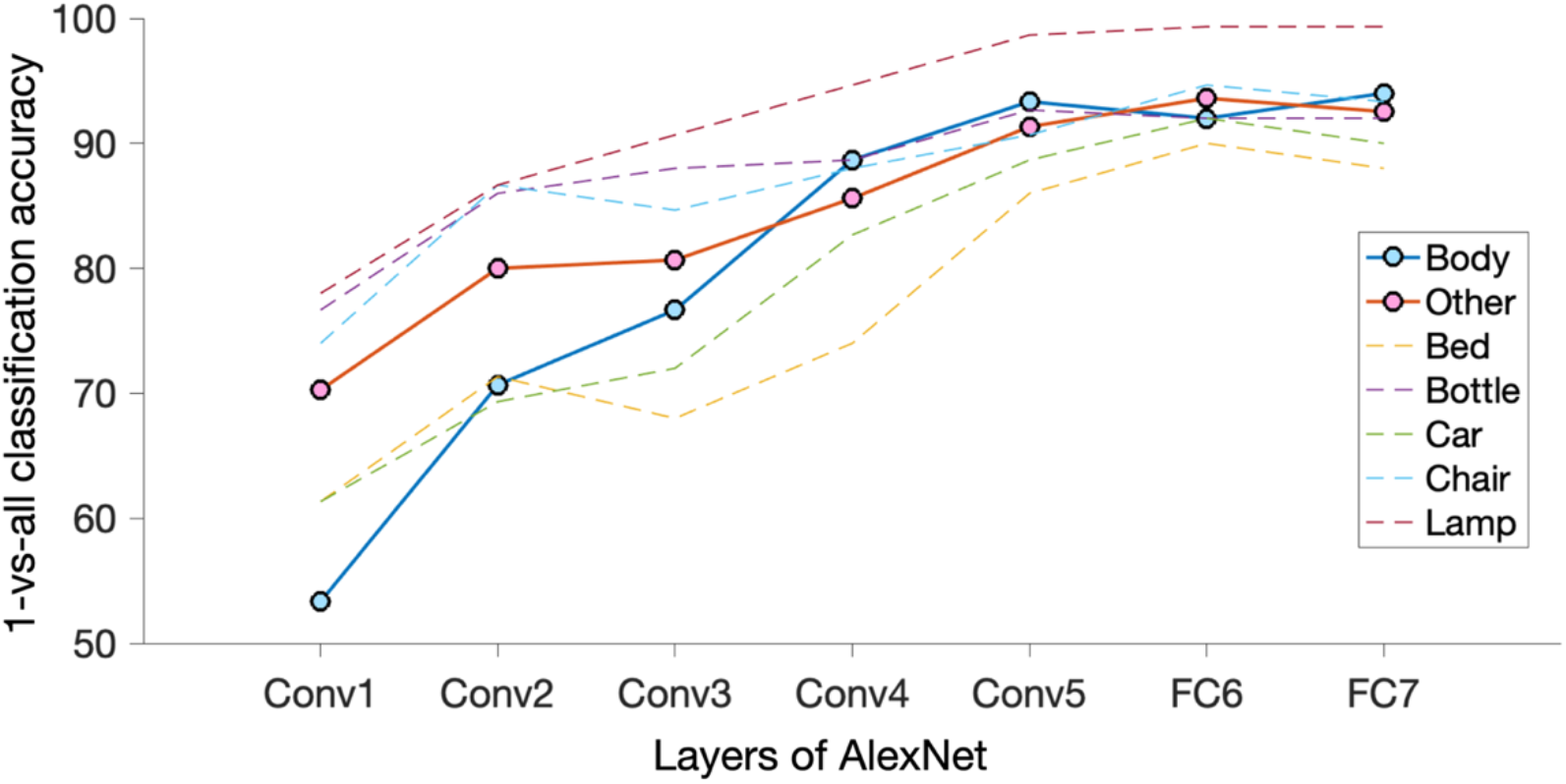
Hierarchical image-based discriminability of the exemplars used in the fMRI experiment. One-vs-all classifiers were trained for each of the six categories, on the output activations of each layer of a convolutional neural network trained for object recognition (AlexNet). 10-fold crossvalidation accuracies are shown for all the objects in addition to the average accuracies for the non-body objects (termed ‘Other’). Discriminability based on low-level features (corresponding to the early layers of the AlexNet) was, if anything, lower for the human bodies than for the other objects. Therefore, it is unlikely that the body-selective fMRI results reflect a distinct low-level property of bodies. ‘Conv’ refers to the convolutional layers of AlexNet and ‘FC’ refers to the fully-connected layers.

## Discussion

Across multiple analyses, we found convincing evidence that attention to human bodies enhanced visual cortex responses selective to bodies presented at task-irrelevant locations. This modulation reflected response gain rather than a generic bias, and could not be explained by low-level feature similarity of bodies. These results indicate that spatially-global attentional modulation – a hallmark of feature-based attention – can be found for features diagnostic of the presence of the human body.

The attentional effects observed here for body silhouettes are unlikely to reflect attention to low-level features such as orientation or color, for several reasons. First, we included a relatively large number of object categories in the experiment to ensure that participants could not detect objects based on low-level features, as these were shared with other categories (e.g., bottles were vertical, similar to bodies). Second, we presented object silhouettes instead of photographs to avoid possible low-level differences between categories in texture or color. Third, the image-based discriminability for each category, established using a convolutional neural network (CNN), indicated that bodies were difficult to discriminate from other categories based on low-level features encoded in the early layers of the CNN. Finally, the fMRI results showed attentional modulation in object-selective cortex (LOC) and body-selective EBA/FBA, but not early visual cortex (EVC), indicating an attentional modulation at a higher level of visual processing.

Our results are in line with the feature similarity gain modulation model (FSGM; Maunsell & Treue, 2006) by showing that feature-based attention enhanced the response to the voxels ‘ preferred stimuli. Specifically, attention to bodies made the response pattern evoked by task-irrelevant bodies more similar to prototypical body response patterns established in a separate baseline experiment. Furthermore, these attention effects were strongest in body-selective voxels of LOC. Finally, reliable univariate attention effects were observed in independently-defined body-selective regions (EBA/FBA). It should be noted that we did not find evidence that responses to the other categories were suppressed, as proposed by FSGM. However, the response to other categories was low and any suppression (posited to be smaller in magnitude than enhancement by FSGM) might not be observable in this case.

The finding of spatially-global modulation for human bodies adds to previous evidence for global modulation for faces. Specifically, in one study, peripherally presented and task-irrelevant faces evoked a stronger face-selective N170 electro-encephalography (EEG) response when participants attended to faces than to houses (Störmer et al., 2019). Furthermore, in fMRI, responses to peripheral faces in the face-selective fusiform face area (FFA) were more strongly modulated by the task-set of the participants (i.e., whether or not they focused on faces) than by spatial attention (Reddy et al., 2007). Together with the current findings, these results provide evidence for spatially-global attentional modulation for bodies and faces, two socially relevant categories that are selectively represented in the visual cortex (Downing et al., 2006; Kanwisher, 2010).

While these results suggest that bodies and faces may be special – reflecting their unique social and biological significance – we do not rule out that spatially-global attentional modulation may also exist for other highly-familiar object categories. For example, behavioral studies showed that animals and vehicles could be detected in the near-absence of spatial attention (Li et al., 2002; but see Cohen et al., 2011), with category-based attention facilitating object detection independently of spatial attention (Stein and Peelen, 2017). Indeed, based on the overlap in human and animal features in detection tasks (Evans and Treisman, 2005), it is plausible that our results would generalize to other animals, particularly those that activate body-selective regions (Downing et al., 2006). Similarly, extensive experience with particular objects may drive selective neural tuning (Gauthier and Logothetis, 2000; McGugin et al., 2012; Frank et al., 2014) and give rise to similar behavioral advantages as those observed for bodies (Hershler and Hochstein, 2009; Golan et al., 2014; Reeder et al., 2016; Stein et al., 2016).

Taking everything together, the evidence suggests that features that are diagnostic of bodies meet many of the previously proposed criteria for basic features: showing spatially-global attentional modulation (Maunsell and Treue, 2006), being processed “early, automatically, and in parallel across the visual field” (Treisman and Gelade, 1980), and being represented selectively in the visual system (Treisman, 2006). Indeed, Treisman (2006) proposed that the feature detectors of the feature integration theory are not necessarily limited to low-level features such as orientation and color. Raising the possibility that there may be animal feature detectors, Treisman (2006) noted that animal features may not necessarily be more complex for the visual system than colors, line orientations, or direction of motion. By providing evidence for spatially-global attentional modulation for human bodies, our results support this proposal.

Our findings raise the question of what features are attended when attention is directed to bodies. Addressing this question for animals, Treisman (2006) suggested that: “participants may be set to sense, in parallel, a highly overlearned vocabulary of features that characterize a particular semantic category.” One possibility is thus that attention to bodies is mediated by attention to a set of mid-level features that are diagnostic of human bodies (Ullman et al., 2002; Reeder and Peelen, 2013). Alternatively, attention may be directed to holistic representations of body shape (Reed et al., 2003; Stein et al., 2012). Future studies could test these alternatives by measuring global attentional modulation for various body-related features, body parts, and inverted bodies at the task-irrelevant location while participants attend to bodies at the task-relevant locations (Reeder and Peelen, 2013).

To conclude, the current results provide evidence for spatially-global attentional modulation for human bodies in high-level visual cortex, linking this modulation to body-selective representations in univariate and multivariate analyses. Combining these results with previous behavioral and neuroimaging studies, we propose that bodies may be processed as basic features, supporting the rapid and parallel detection of conspecifics in our environment even outside the focus of spatial attention.

## Acknowledgements

This project has received funding from the European Research Council (ERC) under the European Union’s Horizon 2020 research and innovation programme (grant agreement No. 725970).

